# *Culex erythrothorax*: Insecticide Susceptibility and Mosquito Control

**DOI:** 10.1101/2020.01.27.921007

**Authors:** AT Esterly, D Alemayehu, B Rusmisel, J Busam, TL Shelton, T Sebay, N Zahiri, JW Huston, RJ Clausnitzer, EJ Haas-Stapleton

**Affiliations:** Alameda County Mosquito Abatement District, 23187 Connecticut Street, Hayward, CA 94545; San Mateo County Mosquito and Vector Control District, 1351 Rollins Road, Burlingame, CA 94010; Currently at Santa Clara County Vector Control District, 1580 Berger Drive, San Jose, CA 95112

## Abstract

*Culex erythrothorax* Dyar is a West Nile virus (WNV) vector that breeds in wetland habitats with emergent vegetation. Urbanization and recreational activities near wetlands place humans and birds in close proximity to *Cx. erythrothorax*, increasing the risk of WNV transmission. High adult *Cx. erythrothorax* abundance peaked in a marsh bordering the San Francisco Bay of California (USA) during the first 3 hours after sunset (5527 ± 4070 mosquitoes / trap night) during which adult mosquito control efforts are likely most effective. These adult mosquitoes were highly sensitive to permethrin, resmethrin, naled, and etofenprox insecticides when tested in a bottle bioassay (LC50 = 0.35, 0.68, 0.71, and 4.1 µg/bottle, respectively). The synergist piperonyl butoxide increased the sensitivity of the mosquitoes to a low concentration of permethrin (0.5 µg/bottle) while also reducing knock down time, but did not have similar effects with a higher permethrin concentration (2.0 µg/bottle). Biochemical enzyme assays measuring bulk mixed-function oxidase, alpha-esterase, or beta-esterase activities in mosquito homogenates were significantly higher in *Cx. erythrothorax* relative to a strain of *Culex pipiens* that is sensitive to permethrin. Larvicide that was applied to the site had limited impact on reducing the abundance of adult *Cx. erythrothorax*. Subsequent removal of emergent vegetation in concert with reduced daily temperature in the environment substantially reduced *Cx. erythrothorax* abundance. Land managers who have a need to control *Cx. erythrothorax* in wetlands should consider vegetation removal over applying larvicide, while vector control agencies are likely to successfully control viremic *Cx. erythrothorax* that enter nearby neighborhoods by applying insecticides that target the adult stage of the mosquito within 3 h of sunset.

## Introduction

*Culex erythrothorax* Dyar, commonly known as the tule mosquito, is endemic to the western southwestern states of the US [1]. The larvae breed in heavily vegetated regions of shallow ponds and can be highly abundant in marsh habitats that contain dense clusters of *Schoenoplectus spp* (common tule), *Typha spp*. (bulrush), or *Myriophyllum aquaticum* (parrot feather) [2–4]. Unlike many species of mosquito, adult *Cx. erythrothorax* do not disperse distantly from where they emerge [3, 5, 6]. The time of host-seeking is well documented for *Culex tarsalis* Coquillett, another mosquito species found marsh habitats, which is reported to be most active 1 – 4 h after sunset [7, 8]. The time of day that *Cx. erythrothorax* is most likely to seek a blood meal and would be best controlled by insecticides that target the adult mosquito is not reported. Larvicide applications to constructed marsh habitats can markedly reduce the abundance of adult *Cx. erythrothorax* [9]. However, the duration of reduction is short, likely because adult mosquitoes immigrate from nearby sites that are not treated [5] and dense aquatic vegetation may limit the penetration of larvicides into the water column.

Adult female *Cx. erythrothorax* are aggressive biters that take blood meals from mammals and birds [6, 10]. West Nile virus (WNV) is an arbovirus that can be transmitted from infected birds or tree squirrels (*Sciurus spp*.) to humans via biting *Cx. erythrothorax* [11]. While about 80 % of WNV infections in humans are apparently asymptomatic, serious neuroinvasive disease develops in less than 1 % of infected persons [12]. The greatest risk for human exposure to WNV is thought to come from biting *Culex pipiens* and *Culex quinquefasciatus*. collected in a marsh habitat that abuts a suburban landscape contained human blood relative to the *Cx. quinquefasciatus* that were collected in the same traps [5] suggesting that the risk of human exposure to WNV via *Cx. erythrothorax* may increase as people seek to reside near and recreate in marsh habitats. Moreover, *Cx. erythrothorax* can maintain the transmission of WNV among birds in marsh habitats, with cooccurring *Cx. tarsalis* transmitting the virus to humans. When larvicides are ineffective in controlling the breeding of *Cx. erythrothorax* larvae, insecticides that target adult mosquitoes may be employed to control breeding and interrupt arbovirus transmission cycles.

Pyrethroid-based and organophosphate insecticides are used by public health agencies to control adult mosquitoes [13]. Pyrethroids are synthetic analogs of natural pyrethrin insecticides, which are typically extracted from the flowers of *Chrysanthemum cinerariifolium* [14], and are the most common active ingredient in household insecticides. The neurotoxic mode of action for pyrethroids is to prevent the transition of the voltage-gated sodium channel (VGSC) from an activated to inactivated state, resulting in continuously depolarized neuronal membranes and paralysis or death of the insect [15, 16]. Organophosphate insecticides, including naled and malathion, act by phosphorylating esterases to inhibit enzymes such as acetylcholinesterase [17, 18]. The loss of active acetylcholinesterase at nerve endings reduces the rate of acetylcholine neurotransmitter hydrolysis, leading to the overstimulation of nicotinic acetylcholine receptors on the surface of neurons and death of the insect [19].

Insecticide resistance in mosquitoes is mediated by genetic changes that reduce insecticide binding to its molecular target, the upregulation of detoxification pathways that reduce the quantity of active insecticides in the mosquito and behavioral changes that reduce the likelihood of mosquitoes encountering insecticides, among others [20, 21]. Several point mutations have been identified in the gene encoding VGSC that are associated with pyrethroid resistance. Most notable are mutations in the *kdr* and *super-kdr* loci of *VGSC* that result in amino acid substitutions at positions 1014 and 918, respectively [22], that likely prevent the insecticide from interacting with VGSC [23]. The upregulation of detoxification pathways results in an increased quantity or activity of enzymes that hydrolyze insecticides within the insect, rendering them inactive and unable to kill the exposed insect. Herein, we examined the activity of alpha- and beta-esterase, glutathione S-transferase (GST), mixed-function oxidase (MFO), and insensitive acetylcholine esterase enzymes in mosquitoes that had not been exposed to insecticides. Esterases are a large family of enzymes that hydrolyze ester bonds within insecticides [24]. MFO transfer reduced oxygen to insecticides while GST conjugate glutathione to insecticides, thereby increasing their solubility in water and rate of excretion from the insect [25–27]. Piperonyl butoxide (PBO) is a synergist that can be included with insecticides to inhibit MFO and increase the efficacy of insecticides [28].

We found that similar to *Cx. tarsalis*, adult *Cx. erythrothorax* sought blood meals in a freshwater marsh habitat that abuts a suburban landscape 1 – 3 h after sunset, with a smaller peak of activity in the hour just before sunrise. The present study describes the insecticide susceptibility of adult female *Cx. erythrothorax* to permethrin, resmethrin, etofenprox and naled, assesses the activity of insecticide detoxifying enzymes in this WNV vector, and demonstrates that removing emergent vegetation from *Cx. erythrothorax* breeding sites should be considered over larvicide applications to reduce mosquito abundance.

## Materials and Methods

### Mosquito Collection

Adult *Culex erythrothorax* mosquitoes were collected over night at the Hayward Marsh, a 0.13 km^2^ freshwater marsh that abuts the San Francisco Bay, CA USA (GPS coordinates: 37.629986, −122.141174) using Encephalitis Vector Survey traps (EVS; BioQuip, Rancho Dominguez, CA) or a Collection Bottle Rotator Trap (CBRT; John W. Hock Company, Gainesville, FL) that were baited with dry ice. EVS traps were placed overnight and the collected mosquitoes enumerated and identified to species using a dissection microscope. Timed mosquito collections over 24 h periods were made using the CBRT that was programed to rotate collection chambers every 3 h. Adult *Cx. tarsalis* were collected in a CBRT that was placed near Bair Island Ecological Reserve, CA USA (GPS coordinates: 37.501533, −122.216144). Mosquitoes that were collected for adult CDC bottle bioassays (BBA) or enzyme activity assays were transported in a humidified chamber and transferred to a nylon mesh chamber (24 x 14 x 13 cm) prior to use. A strain of *Cx. pipiens* mosquitoes that is susceptible to permethrin (strain SM-S1; *Cx. pipiens*^SM-S1^) was reared in an insectary using standard methods. Meteorological data was obtained from a weather station that was located 1.3 km east and 3.2 km north of the Hayward Marsh using the US National Centers for Environmental Information database (www.ncdc.noaa.gov/cdo-web; [29]). Cumulative degree-days (DD) for each week was calculated as described previously for *Culex* mosquitoes [30] by comparing the daily average temperature to a baseline of 10°C (See Equation 1). If the DD calculation resulted in a negative value, zero was used instead.

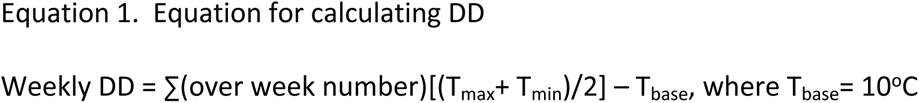

### Adult bottle bioassays

Timed knock-down BBA were conducted to evaluate the resistance of adult mosquitoes to insecticides, as previously described [1]. Briefly, the inside of clear 250 ml graduated media bottles (DWK Life Sciences LLC, Millville, NJ) were evenly coated with technical grade insecticide (permethrin, resmethrin, etofenprox, or naled; Chem Service, West Chester, PA) that was diluted in acetone. PBO (Chem Service, West Chester, PA) was evaluated in combination with permethrin. Control bottles contained only the diluent or diluent with PBO. The diluent was evaporated from the interior of the bottles at room temperature using a gentle steam of nitrogen gas. Adult female mosquitoes were transferred to the bottles (n > 25 mosquitoes per bottle), and the number of dead or knocked down mosquitoes were recorded every 15 min for at least 90 min. A mosquito was recorded as dead or knocked down if it could not stand unaided when the bottle was gently rotated; otherwise, the mosquito was counted as alive.

### Enzyme activity assays

Individual mosquitoes were placed a microcentrifuge tube that contained a 5 mm glass bead, homogenized in potassium phosphate buffer chilled to 4°C using a Bead Mill 24 (Fisher Scientific, Waltham, MA) for 25 s at a speed setting of 4.0, and the homogenate clarified using centrifugation (3 min, 10,000 x g). Enzyme assays were conducted as described previously to evaluate the activity of MFO [31],GST [32], alpha-esterase [33], beta-esterase [33], and for the insensitive acetylcholinesterase assay [34]. Protein content of each mosquito homogenate was determined to correct for differences in mosquito size using a Pierce BCA protein assay kit, as described by the manufacturer (Thermo Scientific, Waltham, MA). Absorbances were measured using an Epoch Microplate Spectrophotometer (BioTek Instruments, Winooski, VT).

### Larvicide application and vegetation removal for mosquito control

Approximately 100 kg of Vectolex FG (Clarke, St. Charles IL), VectoMAX FG (Valent Bio Sciences, Libertyville, IL), or Vectobac G (Valent Bio Sciences, Libertyville, IL) larvicide was applied at the Hayward Marsh near the edges of emergent bulrush vegetation every 1 – 3 weeks at a rate of 22 kg / hectare using a Mist Duster MD 155DX powered backpack blower (Maruyama US, Fort Worth, TX) by walking the perimeter of the marsh or by boat to access the emergent vegetation that was not adjacent to the shoreline. One product was used during each application week and the products were rotated in the order indicated above. Mosquito abundance was evaluated every 1 – 3 weeks at Hayward Marsh using 3 – 10 EVS traps that were baited with dry ice, as described previously. The quantity of emergent vegetation that was removed from the site between October 2015 and March 2017 was estimated by drawing bounding boxes around the regions using Google Earth Pro (version 7.3.2.5776).

### Statistical Methods

Data was plotted and analyzed using Prism software (version 8.3.0; GraphPad Software, San Diego, CA). Each insecticide concentration was assessed in triplicate BBA for each species and the mean ± the standard error of the mean (SEM) reported. The lethal concentration of pesticide that knocked down or killed 50% of the mosquitoes (LC50) in BBA was calculated from the equation of the line that was generated from a linear regression of the dose-mortality data for each pairing of species and pesticide. The resistance ratio (RR) was calculated by dividing the LC50 values of the field-collected *Cx. erythrothorax* by the LC50 value of the SM-S1 strain of *Cx. pipiens*. The slope and intercepts of linear regressions were evaluated with an analysis of covariance, and 0.05 set as the threshold of significance. Enzyme activity assays were conducted in triplicate for each mosquito homogenate (n = 10 mosquitoes per species). All t tests were two tailed with the threshold of significance set at 0.05.

## Results and Discussion

### Hourly adult activity times of Cx. erythrothorax relative to Cx. Tarsalis in marsh habitats

The Collection Bottle Rotator Trap (CBRT) was operated over three consecutive 24 h periods and captured 10041 ± 5332 *Cx. erythrothorax* and 267 ± 74 *Cx. tarsalis* per day (Fig 1). The proportion of the total mosquitoes for each species that were collected in CBRT peaked at 1 – 3 h after sunset for *Cx. erythrothorax* (5527 ± 4070 adult female *Cx. erythrothorax*; Fig 1), while host-seeking *Cx. tarsalis* were most abundant 3 – 6 h after sunset (83 ± 30 adult female *Cx. tarsalis*; Fig 1). Blood meal-seeking, as measured by catch in the CBRT, decreased markedly for both mosquito species during and subsequent to the 12 – 15 h collection period, which correlated with the collection period immediately after sunrise (Fig 1). Nocturnal host seeking behavior by female *Cx. erythrothorax* was similar to that of *Cx. tarsalis*, *Cx. pipiens* and *Anopheles gambiae* [7, 35, 36]. *Culex erythrothorax* grow in marsh habitats that contain bulrush or tule vegetation where waterfowl can be abundant. The host seeking activity of *Cx. erythrothorax* correlates with when waterfowl roost during the nighttime.

**Fig 1.**
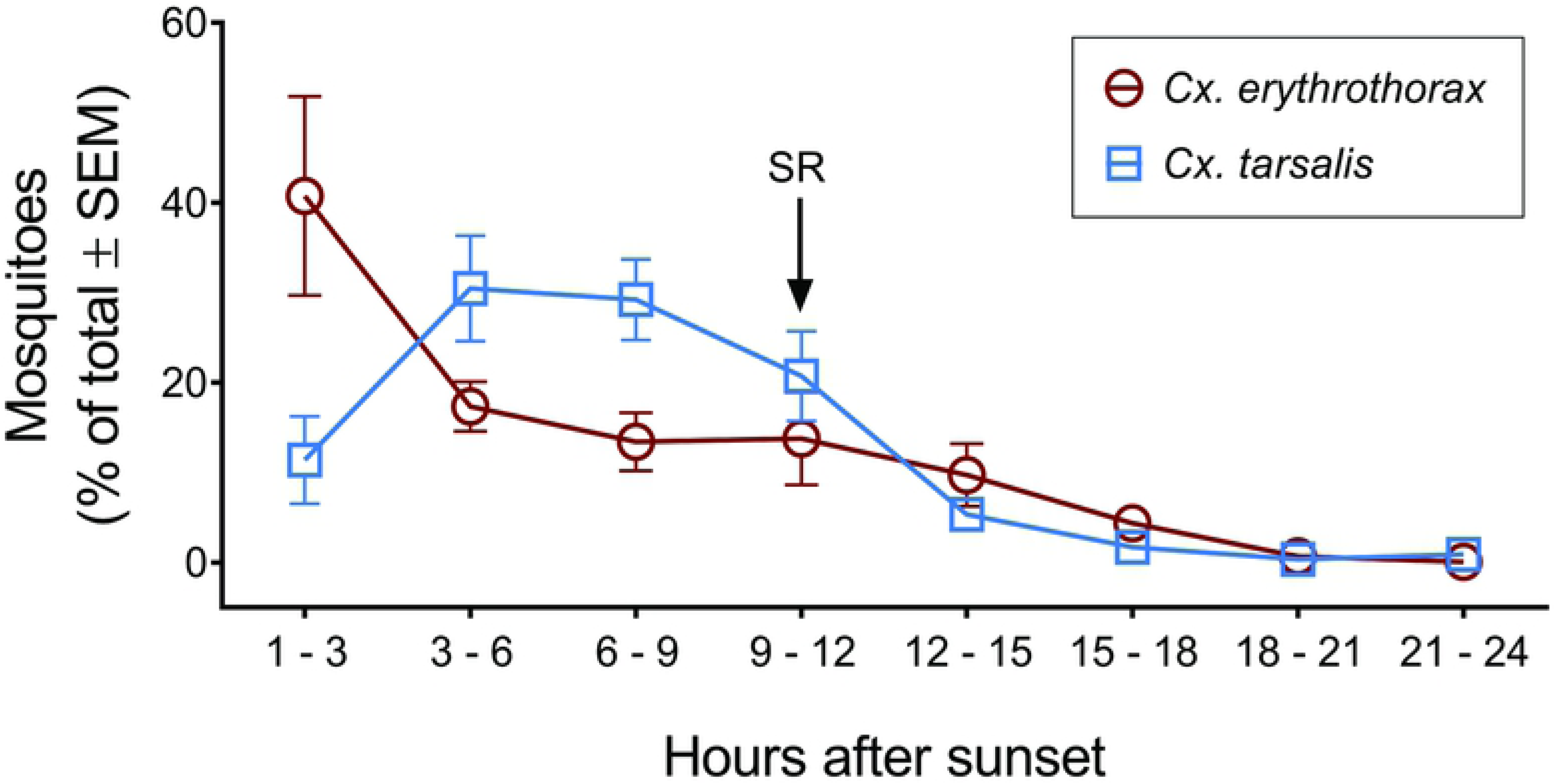
Hourly abundance of *Cx. erythrothorax* and *Cx. tarsalis* in a marsh habitat measured using a collection bottle rotator trap. The average proportion of mosquitoes that were collected during 3 h periods after sunset for three days is shown for *Cx. erythrothorax* in red and *Cx. tarsalis* in blue. SR indicates the time period during which sunrise occurred.

### Bioassays evaluating insecticide resistance

Assessing insecticide resistance in mosquito populations that are collected from habitats that are not degraded may provide the opportunity to establish a regional baseline for insecticide resistance in mosquitoes. A CDC bottle bioassay (BBA) was used to assess the susceptibility of *Cx. erythrothorax* that were collected from a marsh habitat that supports diverse wildlife to permethrin, an insecticide that is widely used to control mosquitoes and other insect pests. Mosquito knock down or mortality in bottles that contained only vehicle or PBO was not greater than 3% (not shown). The lethal concentration that knocked down 50% of the mosquitoes after a 90 min exposure to permethrin was 9-fold lower for *Cx. erythrothorax* relative to *Cx. pipiens*^SM-S1^ indicating that the former was highly susceptible to permethrin (Fig 2; LC50 of 0.35 and 3.1 µg / bottle, respectively; resistance ratio (RR) of 0.11). There was no significant difference in the slopes of the linear regression lines for both species exposed to permethrin, suggesting that their biological responses to permethrin were similar (F = 2.492, DFn = 1, DFd = 5, P = 0.1753). However, the y-intercept of the lines were significantly different, indicating that the field-caught *Cx. erythrothorax* were intrinsically more sensitive to permethrin relative to *Cx. pipiens*^SM-S1^ (y-intercept of 68.3 ± 13.3 and 101.0 ± 15.0, respectively; F = 10.73, DFn = 1, DFd = 6, P = 0.0169).

**Fig 2.**
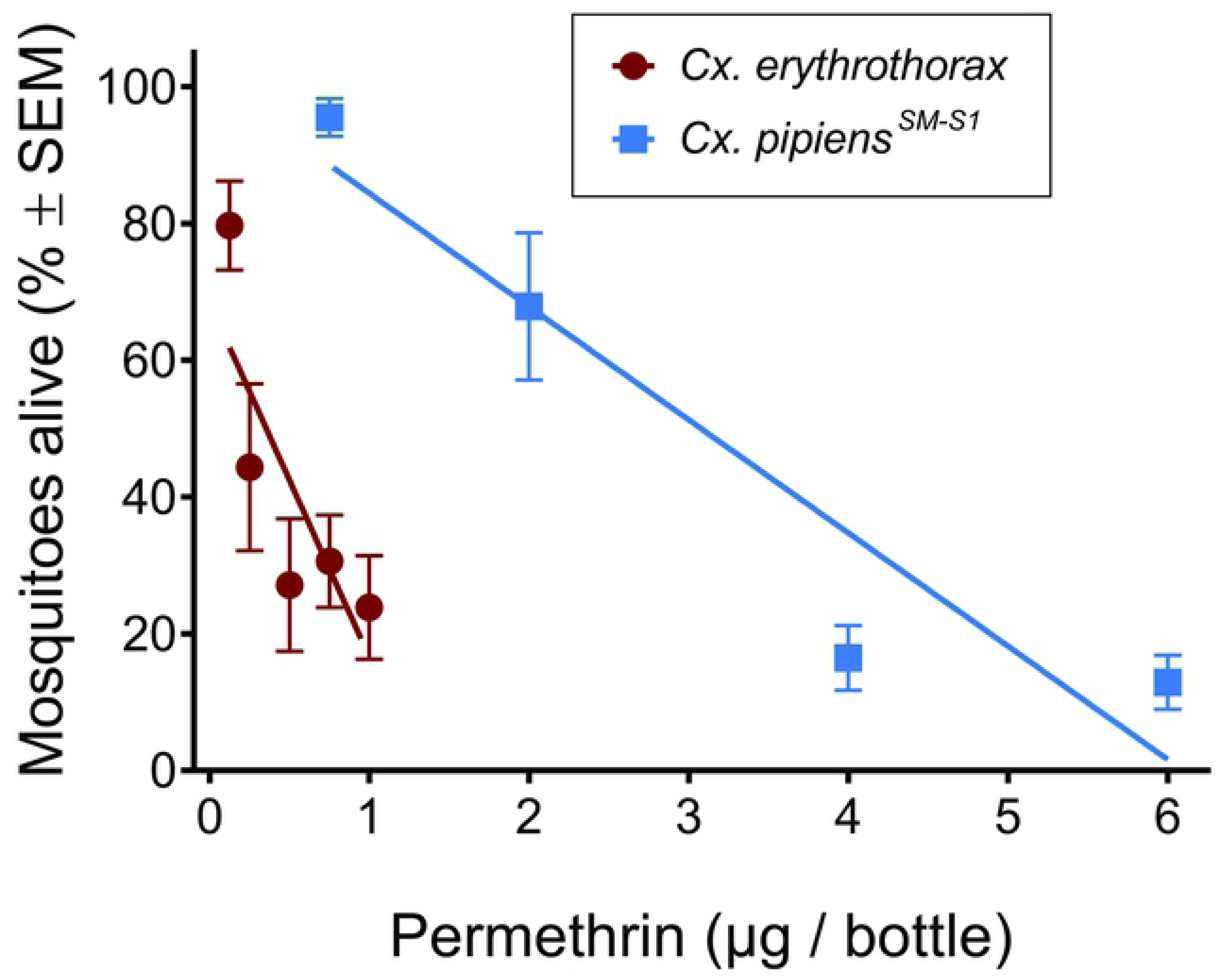
Susceptibility of *Cx. erythrothorax* and *Cx. pipiens*^SM-S1^ to permethrin. The average proportion of mosquitoes that remained alive after exposure to permethrin for 90 min in a BBA with *Cx. erythrothorax* in red and *Cx. pipiens*^SM-S1^ in blue.

Exposing adult *Cx. erythrothorax* for 90 min to 5 µg or 20 µg of PBO per bottle in the absence of permethrin did not significantly affect mosquito knockdown relative to mosquitoes in bottles that lacked PBO (Fig 3; unpaired t test, P = 0.5048 and 0.7452, respectively). Inclusion of 5 µg or 20 µg of PBO to bottles containing 0.5 µg of permethrin significantly reduced the proportion of live *Cx. erythrothorax* after 90 min relative to mosquitoes in bottles lacking PBO, although the difference was numerically small (Fig 3; unpaired t tests, P = 0.0346 and 0.0257, respectively; 1.4 – 1.6 fold reduction). Neither of the PBO concentrations that were evaluated significantly affected survivorship of *Cx. erythrothorax* that were exposed to bottles containing 2.0 µg of permethrin (Fig 3; unpaired t tests, P = 0.5478 and 0.1321, respectively).

**Fig 3.**
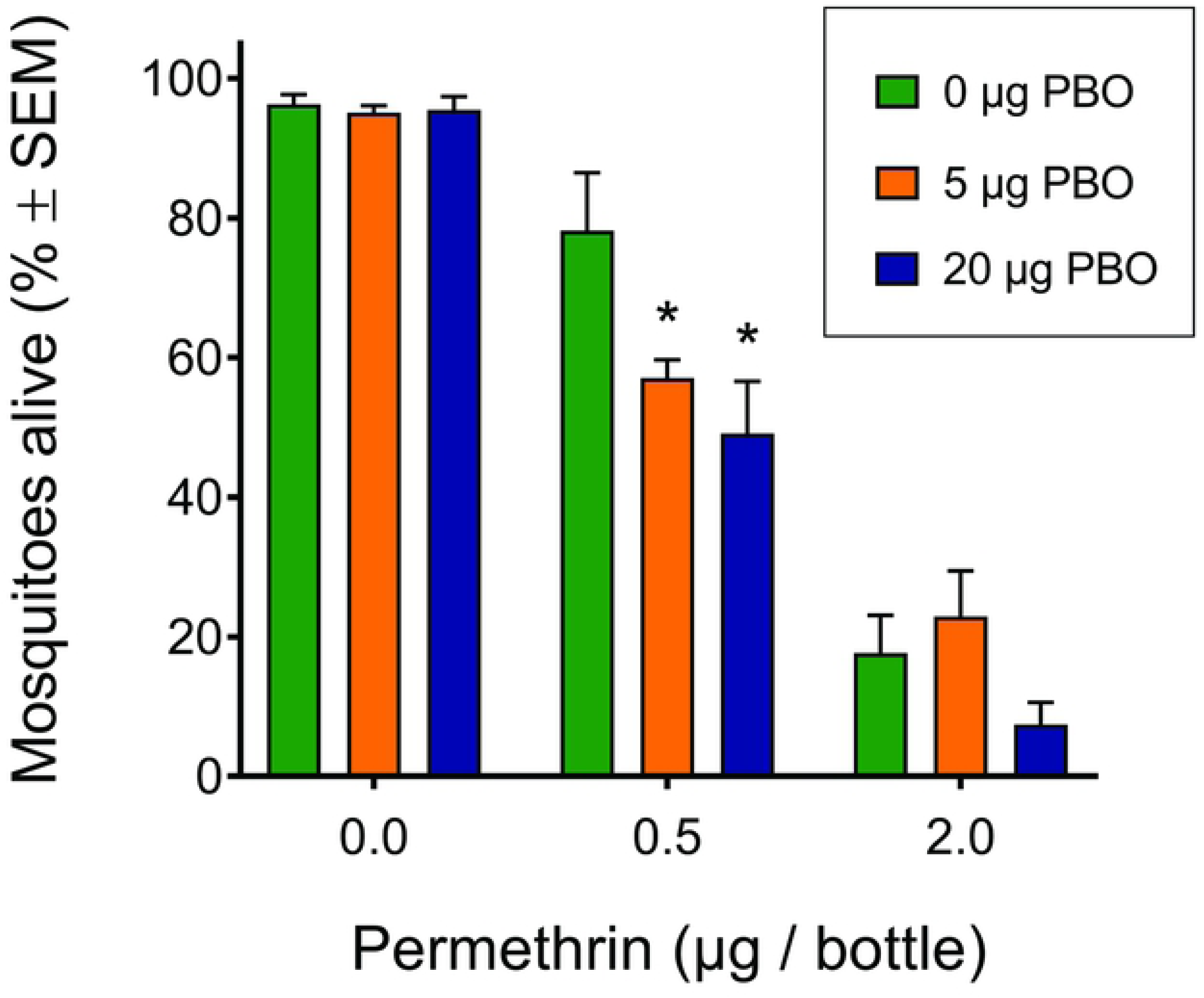
The synergist PBO decreases the survivorship of *Cx. erythrothorax* that were exposed to permethrin. The average proportion of mosquitoes that remained alive after exposure to 0.5 or 2.0 µg of permethrin with or without 5 or 20 µg of PBO. Asterisks (*) indicate a significant reduction in the proportion of living mosquitoes relative to the 0.5 µg permethrin with 0 µg PBO treatment.

The proportion of *Cx. erythrothorax* that were not knocked down 15 min to 90 min after exposure to bottles containing 0.5 µg of permethrin with or without 5 µg of PBO decreased linearly and the rate of knockdown was not significantly affected by the inclusion of PBO (Fig 4A, R^2^ of 0.8483 and 0.9356; F = 4.053, DFn = 1, DFd = 10, P = 0.0718). However, the y-intercept of the regression lines for *Cx. erythrothorax* exposed to 0.5 µg of permethrin with or without 5 µg of PBO were significantly different demonstrating that PBO affected mosquito knockdown (Fig 4A; F = 7.858, DFn = 1, DFd = 11, P = 0.0172). The time to knock down 50 % of the mosquitoes that were exposed to 0.5 µg of permethrin (i.e. KDT50) was 104 min (Y = −0.5186*X + 103.9; R^2^ = 0.9356) and 160 min (Y = −0.3403*X + 104.3; R^2^ = 0.8483) for treatments with or without 5 µg of PBO, respectively. This corresponded to a 35 % reduction in KDT50 when 5 µg of PBO was included in the bottle and 43 % reduction with 20 µg of PBO. Including 20 µg of PBO in bottles that contained 0.5 µg of permethrin significantly increased the mortality rate of *Cx. erythrothorax* relative to mosquitoes that were exposed to 0.5 µg of permethrin in the absence of PBO (Fig 4A; KDT50 = 91 min; Y = −0.5774*X + 102.7; R^2^ of 0.9327; F = 6.28, DFn = 1, DFd = 10, P = 0.0311). While the proportion of *Cx. erythrothorax* that survived exposure to 2.0 µg of permethrin also decreased linearly from 15 min to 90 min, the rate of death and mortality was not significantly affected by the inclusion of 5 µg or 20 µg of PBO (Fig 4B; F = 0.766, DFn = 2, DFd = 15, P = 0.4822). The KDT50 values for *Cx. erythrothorax* that were exposed to 2.0 µg of permethrin with or without PBO were lower than the KDT50 values for mosquitoes exposed to 0.5 µg of permethrin (KDT50 for 2.0 µg permethrin tests: 0 µg PBO (48 min; R^2^ = 0.9041; Y = −1.153*X + 105.3); 5 µg PBO (58 min; R^2^ = 0.9265; Y = −0.9989*X + 108.6); 20 µg PBO (43 min; R^2^ = 0.9678; Y = −0.9332*X + 90.59). In aggregate, the results demonstrate that *Cx. erythrothorax* were exquisitely sensitive to permethrin, and that inclusion of 5 or 20 µg of PBO with 0.5 µg permethrin increased the sensitivity of the mosquitoes to permethrin and hastened knockdown.

**Fig 4.**
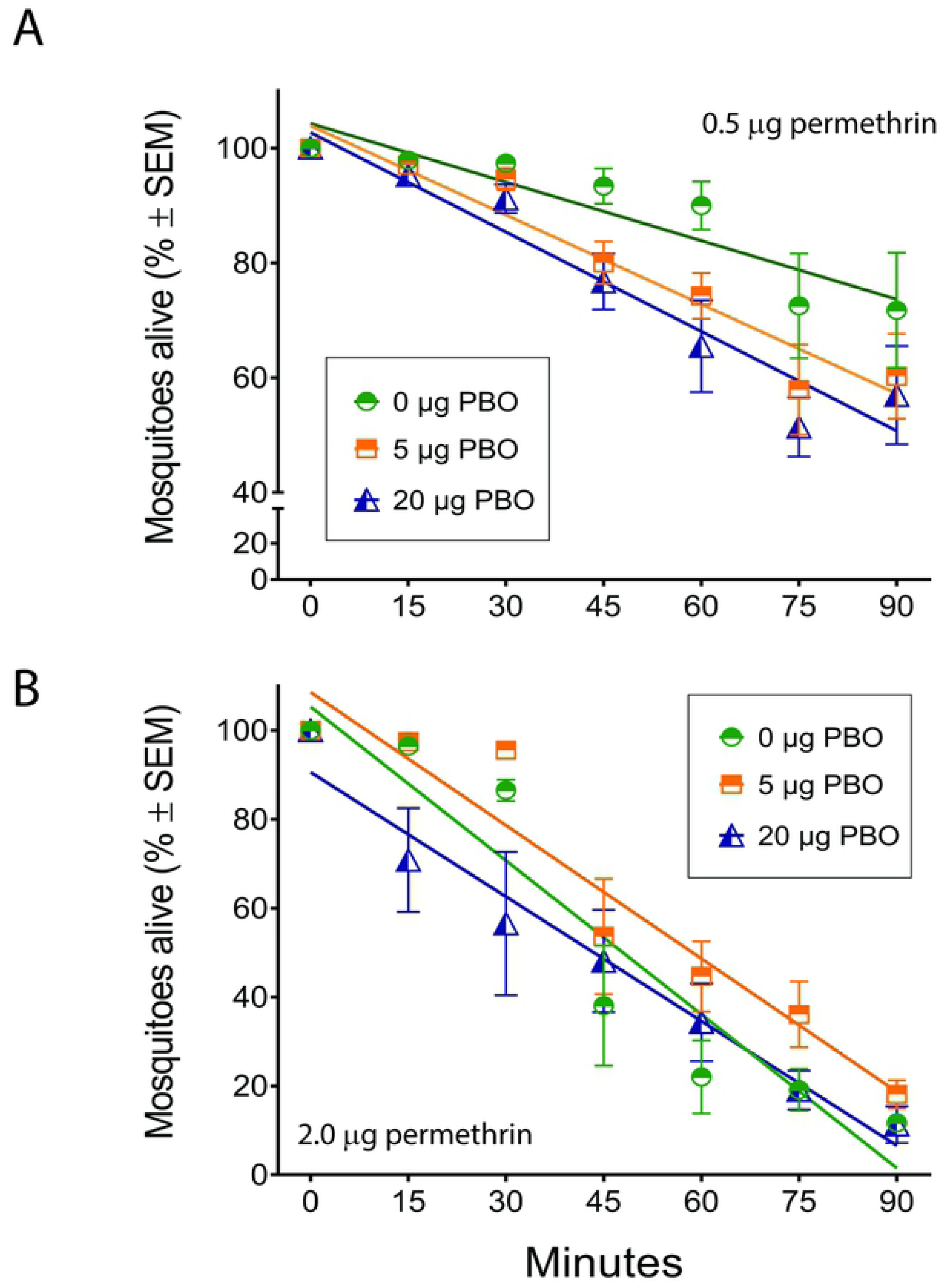
Time that *Cx. erythrothorax* remained alive after exposure to permethrin with or without PBO in a BBA. The average proportion of *Cx. erythrothorax* that remained alive at 15 min intervals after exposure to 0.5 µg of permethrin (4A) or 2.0 µg of permethrin (4B) in the presence of 0, 5 or 20 µg of PBO. Green lines indicate 0 µg of PBO, orange lines 5 µg of PBO, and blue lines 20 µg of PBO.

Adult *Cx. erythrothorax* also displayed high sensitivity to the pyrethroids resmethrin and etofenprox relative to *Cx. pipiens*^SM-S1^ (Fig 5A and, 5B; resmethrin: LC50 = 0.68 µg and 0.24 µg for *Cx. erythrothorax* and *Cx. pipiens*^SM-S1^, respectively with an RR of 2.8; etofenprox: LC50 = 4.1 µg and 8.1 µg for *Cx. erythrothorax* and *Cx. pipiens*^SM-S1^, respectively with an RR of 0.50). Similarly, adult *Cx. erythrothorax* were exquisitely sensitive to naled, an organophosphate insecticide, relative to susceptible strain of *Cx. pipiens*^SM-S1^ (Fig 5C; LC50 = 0.71 µg and 4.0 µg of naled for *Cx. erythrothorax* and *Cx. pipiens*^SM-S1^, respectively with an RR of 0.18).

**Fig 5.**
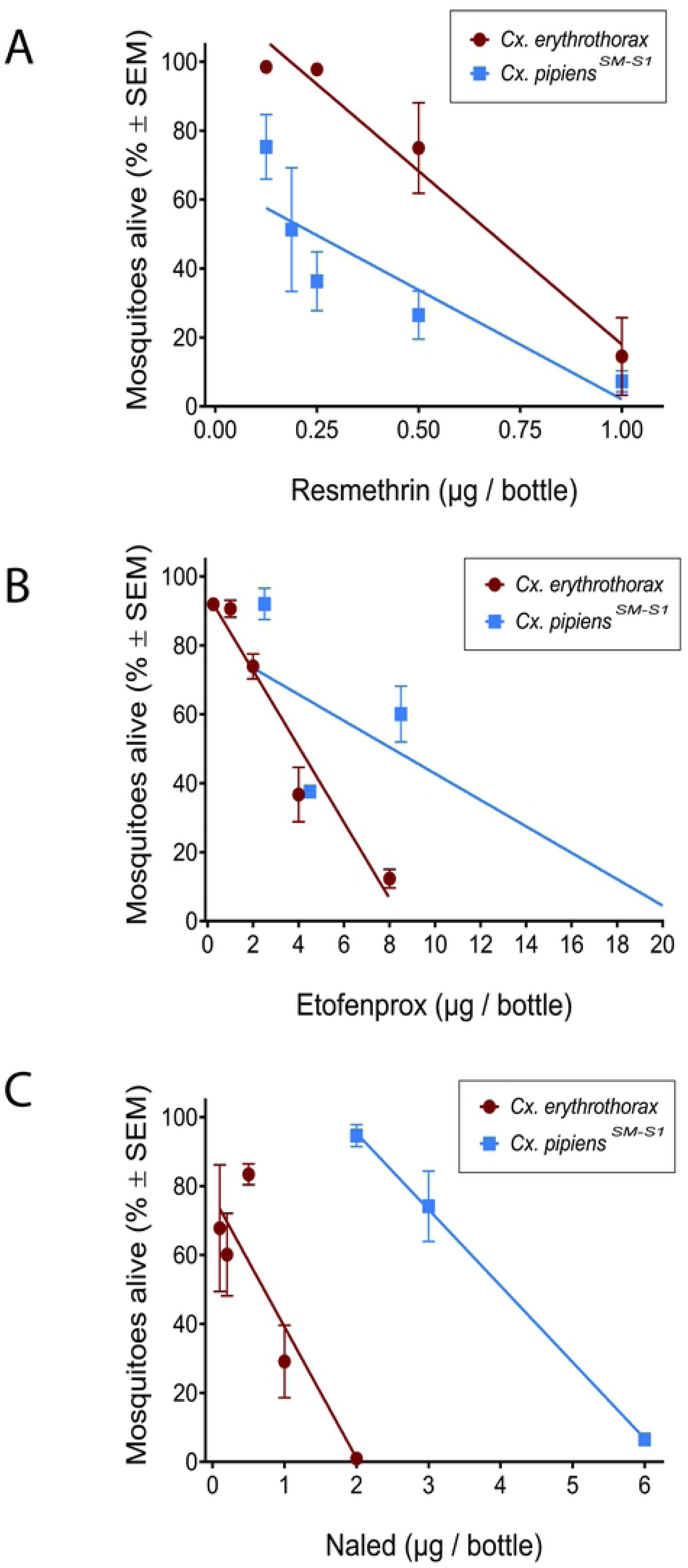
Proportion of mosquitoes alive after exposure for 90 min to resmethrin (A), etofenprox (B) or naled (C) in a BBA. Red lines indicate *Cx. erythrothorax* and blue indicate *Cx. pipiens^SM-S1^*.

As people build residential and commercial properties increasingly closer to wetlands, it is likely that highly abundant mosquitoes, such as *Cx. erythrothorax*, will migrate into these communities seeking a blood meal before returning to the wetland to oviposit. Many bird species that utilize wetland habitats are effective reservoirs for WNV [37]. The sympatric distribution of abundant *Cx. erythrothorax* and marsh birds that are susceptible to WNV with people that live, work and recreate near wetlands may increase the potential for WNV transmission to people. Although pyrethroids should not be applied to areas where surface water is present, such as marsh habitats, permethrin in conjunction with the synergist PBO could be effectively employed by public health agencies to control viremic *Cx. erythrothorax* that enter human communities when seeking blood meals.

### Activity of insecticide metabolizing enzymes

Biochemical assays can be employed to compare the activity of enzymes that detoxify insecticides in field-caught mosquitoes relative to susceptible reference strain. The potential of the field-caught adult *Cx. erythrothorax* to metabolize and inactivate insecticides was evaluated by measuring the activity of detoxifying enzymes in homogenates of individual mosquitoes. There was no significant difference in the activity of GST between *Cx. erythrothorax* and *Cx. pipiens*^SM-S1^, and insensitive acetylcholinesterase was not detected in either species (Fig 6, unpaired t test, P = 0.0932). The *Cx. erythrothorax* displayed significantly higher enzyme activity for MFO, alpha-esterase, and beta-esterase relative to *Cx. pipiens*^SM-S1^ (Fig 6; unpaired t tests, P < 0.0001). Considering that the *Cx. erythrothorax* were more sensitive than *Cx. pipiens*^SM-S1^ to permethrin, etofenprox and naled (Fig 1 and 5), the higher enzyme activity in *Cx. erythrothorax* may be present to metabolize plant phytochemicals or environmental toxins that leach into the water [38]. The KDT50 values for *Cx. erythrothorax* exposed to permethrin in BBA (Fig 4) were higher than what has been reported previously for *Culex spp*. that are resistant to permethrin [39–41]. The increased activity of detoxifying enzymes in *Cx. erythrothorax* (Fig 6) or unmeasured factors such as differences in cuticular structure or chemistry may have contributed to the difference in KDT50 values [42].

**Fig 6.**
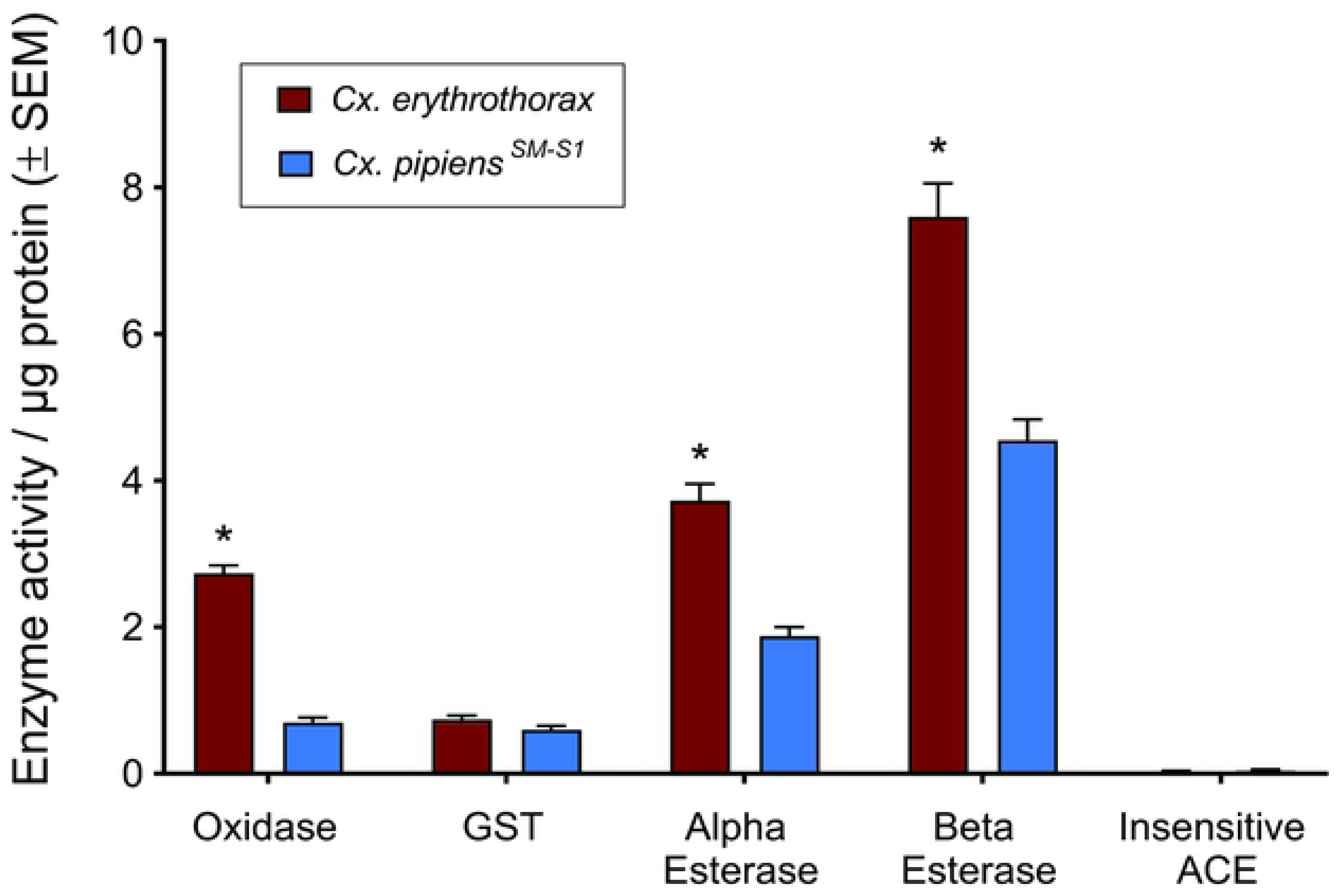
Activity of insecticide detoxifying enzymes normalized to the quantity of mosquito protein. Red bars indicate *Cx. erythrothorax* and blue indicate *Cx. pipiens^SM-S1^*. Asterisks (*) indicate significant differences in the activity of an enzyme between the species.

### Mosquito control: larvicide applications and vegetation removal

Enhanced monitoring at the Hayward Marsh for adult *Cx. erythrothorax* was conducted from week 13 – 52 of 2016 using CO_2_-baited EVS traps (Fig 7). At least 3 EVS traps were placed at the site for each week that abundance was monitored. Over this period, abundance of adult female *Cx. erythrothorax* ranged from 2.7 – 5034 mosquitoes per trap night and averaged 827 ± 156 mosquitoes per trap night. Average weekly wind speeds did not differ substantially over the study period (Fig 7), suggesting that variations in mosquito abundance was not attributed to the capacity of EVS traps to capture mosquitoes or displacement of the mosquitoes from the marsh by wind. Because during week 27 the abundance of adult *Cx. erythrothorax* exceeded 1000 mosquitoes / trap night, approximately 100 kg of larvicide was applied at the site every 1 – 2 weeks from week 27 – 46 of 2016 (Fig 7). Changes in weekly DD tracked with mosquito abundance, with the exception of the period between week 29 – 34 when mosquito abundance was low while DD was high (Fig 7), suggesting that the application of larvicide during this period may have reduced mosquito abundance. Alternatively, the observed reduction in *Cx. erythrothorax* abundance during that period may be due to other environmental or biological factors that could not be assessed with the available data.

**Fig 7.**
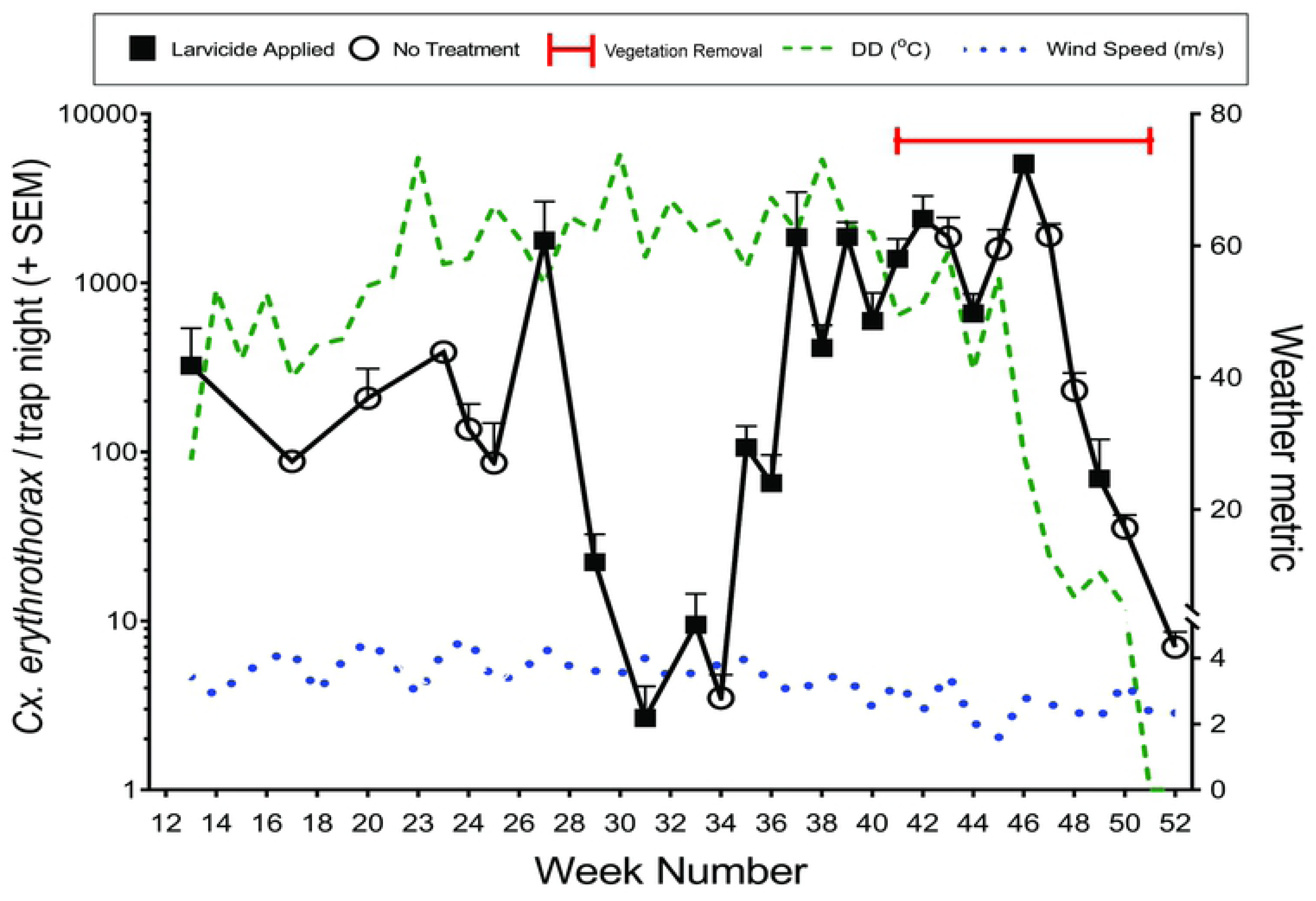
Average weekly abundance of *Cx. erythrothorax* in a marsh habitat relative to wind speed and degree days (DD). Open circles indicate weeks that larvicide was not applied and closed squares indicate weeks that larvicide was applied. The red bar indicates the period during which emergent vegetation was removed from the site.

Although larvicide was applied at the site during weeks 37 – 42, the dense emergent vegetation supported high mosquito abundance (Fig 7; 1421 ± 290 *Cx. erythrothorax* / trap night). Bands of emergent vegetation that are 20 m wide support high *Cx. Erythrothorax* abundance even when larvicide is applied [43]. Because the width of emergent vegetation at the Hayward Marsh exceeded 80 m near the center of the marsh (Fig 8A), the larvicide was unlikely able to penetrate the vegetation and enter the water to impact the mosquito larvae that were *in situ*. To reduce the suitability of the habitat for *Cx. erythrothorax*, approximately 1.5 x 10^4^ m^2^ of emergent vegetation was removed from the marsh during weeks 41 – 51 of 2016, reducing the maximum width of the emergent vegetation to 7 m at the periphery of the march (Fig 8B). The abundance of adult *Cx. erythrothorax* was reduced substantially after the emergent vegetation was reduced (Fig 8A and 8C), however it was coincident with a concomitant reduction in DD (Fig 7). Adult *Cx. erythrothorax* abundance remained relatively low during the two years that followed the reduction of emergent vegetation at the site (77.4 ± 19.3 mosquitoes / trap night). When faced with unacceptably high abundance of *Cx. erythrothorax* in a marsh, land managers may consider forgoing intensive and costly larvicide applications if the width of emergent vegetation is high, and instead focus on removing the vegetation. In doing so, subsequent larvicide applications are more likely to reduce the growth of mosquito larvae while increasing the ability of fish and invertebrates to prey upon the mosquito larvae.

**Fig 8.**
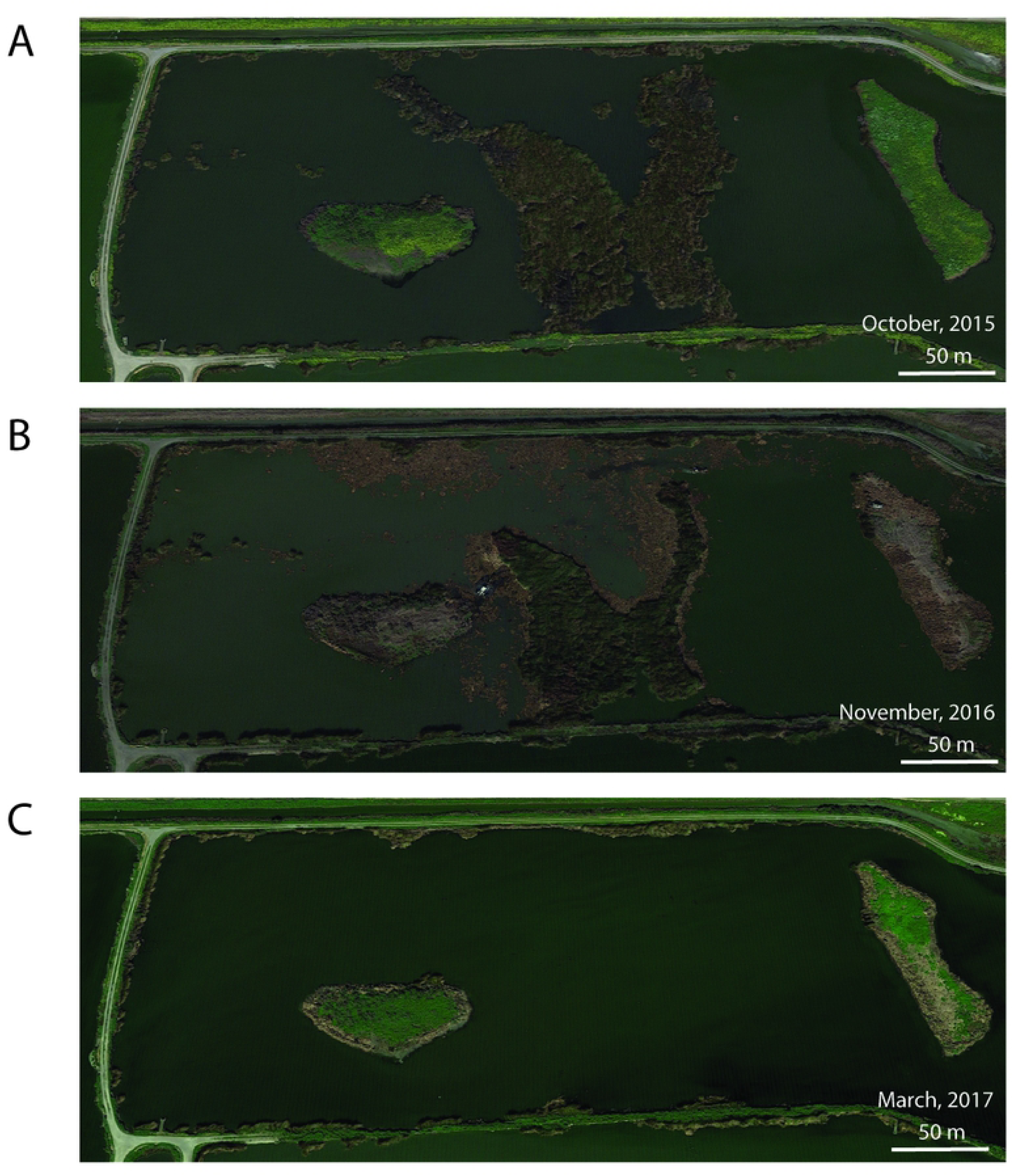
Aerial imagery of the site before (A), during (B) and after (C) the removal of emergent vegetation. Dates in the lower right corner of each image indicate the date that the image was collected. Map data: Google, Maxar Technologies.

## Conclusions

Adult *Cx. erythrothorax* in the present study were highly susceptible to several pyrethroid-based insecticides and naled, an organophosphate insecticide, even though they were collected less than 1 km from an industrial area where insecticides are utilized for structural pest control. The incongruency of low LD_50_ with high KDT50 values potentially points to an unappreciated resistance mechanism in *Cx. erythrothorax* that may provide the mosquitoes sufficient time to escape from where the insecticide was applied and survive. Adult *Cx. erythrothorax* likely become infected with arboviruses while they are in marsh habitats when viremic birds are present. However, the insecticides that were evaluated (which are typically employed by public health agencies to control adult mosquitoes) should not be applied over natural bodies of water. Thus, the use of insecticides that target adult *Cx. erythrothorax* can be utilized only when the mosquitoes enter nearby areas that humans occupy when seeking refuge or a blood meal. The time of day that *Cx. erythrothorax* are most actively seeking a blood meal coincides with that of *Cx. tarsalis*, *Cx. pipiens* and *Cx. quinquefasciatus*, each of which can transmit arboviruses such as WNV to people. Consequently, efforts to control viremic *Cx. erythrothorax* in areas that surround marsh habitats may also be effective against these and other crepuscular *Culex* species. Larvicides that are applied to water are typically highly effective in controlling the larvae of most mosquito species. Although there appeared to be a temporal window in the current study where *Cx. erythrothorax* may have been controlled by the intensive larvicide applications, the impact was short lived, and the effort had a high financial cost. An effective approach for controlling *Cx. erythrothorax* larvae in a marsh with dense emergent vegetation is to remove the dense vegetation [44]. By doing so, the habitat can no longer provide the environmental conditions needed by *Cx. erythrothorax*, and would allow larvicide that is applied to enter the water column where the mosquitoes live. The biomass of a single *Schoenoplectus* plant (*i.e.* bulrush) can increase by 0.5 – 3.3 kg in a single year [45, 46], pointing to the importance of implementing an ongoing vegetation management program in mash habitats that abut urban and suburban areas to keep *Cx. erythrothorax* abundance low.

## Acknowledgements

We thank Mark Taylor with East Bay Regional Park District for providing access to the study site and assistance with larvicide applications. We greatly appreciate Noor Tietze with Santa Clara County Vector Control District for providing comment on the manuscript that improved clarity.

